# Differentiation-dependent chromosomal organization changes in normal myogenic cells are absent in rhabdomyosarcoma cells

**DOI:** 10.1101/2023.05.11.540394

**Authors:** Joe Ibarra, Tyler Hershenhouse, Luay Almassalha, Kyle L. MacQuarrie

## Abstract

Myogenesis, the progression of proliferating skeletal myoblasts to terminally differentiated myotubes, regulates thousands of target genes. Uninterrupted linear arrays of such genes are differentially associated with specific chromosomes, suggesting chromosome specific regulatory roles in myogenesis. Rhabdomyosarcoma (RMS), a tumor of skeletal muscle, shares common features with normal muscle cells. We hypothesized that RMS and myogenic cells possess differences in chromosomal organization related to myogenic gene arrangement. We compared the organizational characteristics of chromosomes 2 and 18, chosen for their difference in myogenic gene arrangement, in cultured RMS cell lines and normal myoblasts and myotubes. We found chromosome-specific differences in organization during normal myogenesis, with increased area occupied and a shift in peripheral localization specifically for chromosome 2. Most strikingly, we found a differentiation-dependent difference in positioning of chromosome 2 relative to the nuclear axis, with preferential positioning along the major nuclear axis present only in myotubes. RMS cells demonstrated no preference for such axial positioning, but induced differentiation through transfection of the pro-myogenic miRNA miR-206 resulted in an increase of major axial positioning of chromosome 2. Our findings identify both a differentiation-dependent, chromosome-specific change in organization in normal myogenesis, and highlight the role of chromosomal spatial organization in myogenic differentiation.

## Introduction

Myogenesis, the development of skeletal muscle, is a tightly regulated and coordinated sequential process that results in cells proceeding from the state of proliferating myoblast to terminally differentiated myotube^1^. Through the feed-forward action of a small number of myogenic regulatory factors^2-4^, hundreds of genes are significantly differentially regulated during the process^5 6^, ultimately resulting in the final post-mitotic, multi-nucleated form of myotubes.

These large-scale expression changes have been shown to be associated with changes in the organization of nuclear space at such levels as regions of the genome^7 8^, the positioning of specific chromosomal areas^9^, and the association between centromeres and nucleoli^10^. Some of these changes have been demonstrated to play causative roles in the regulation of involved genes^7 9^, demonstrating the importance of changes in the organization of the components of the nucleus to the process of myogenesis as a whole.

Rhabdomyosarcoma (RMS) is a tumor of skeletal muscle, and the most common of the pediatric soft tissue sarcomas^11^. RMS possess many similarities to normal myogenic cells: from morphology, to gene expression of the myogenic regulatory factors^12 13^, to the genome-wide binding patterns of myogenic regulatory factors to DNA^5^. Despite those similarities, RMS cells suffer from a differentiation defect that permits them to continue to proliferate, unless induced to differentiate by various mechanisms^5 13-16^, many of which affect components of the normal myogenic pathway. RMS is also notable for the fact that some tumors bear a characteristic PAX-FOXO transcription factor gene fusion, while others do not^17^. Gene fusion status impacts not only multiple facets of tumor cell biology^18-26^, but clinical outcomes as well^27^. Despite their similarity to normal myogenic cells, less is known about what deficits may or may not be present in RMS at the level of higher-order organization, such as chromosomal positioning and genomic organization.

In this study, we utilized cultured cells to investigate the organizational characteristics of two chromosomes – chromosome 2, which has been shown to be enriched for tandem gene arrays (TGAs) of myogenic genes, and chromosome 18, which is not enriched for myogenic TGAs ^9^. By comparing the characteristics of those chromosomes in multiple RMS cell lines, both those with and without the characteristic PAX-FOXO gene fusion, to the characteristics seen in both proliferating and differentiated normal myocytes, we sought to 1) characterize the extent of similarities and differences in chromosomal organization both between the tumor cell lines and tumor and normal cells and 2) identify potential chromosomal level organizational deficits in RMS cells. We found that, while both chromosomes occupied greater nuclear area in all tumor cell lines compared to normal cells, the positioning of chromosome 2 relative to the nuclear periphery was largely preserved in the tumor cells. In contrast, chromosome 18 exhibited a greater frequency of occupying the more peripheral area of the nucleus in tumor compared to normal cells. More strikingly, we found a chromosome- and differentiation-specific difference in positioning relative to the nuclear axis. In differentiated myotubes, chromosome 2 was found positioned close to the major axis of the cell far more frequently than would be expected by chance, while the positioning in myoblasts and RMS cells did not show this predilection. In contrast, chromosome 18 positioning was indistinguishable from what would be expected by chance in all cell types and conditions. Induced differentiation in the RMS cells increased the frequency of chromosome 2 positioning along the major axis of the cell, suggesting a relationship between chromosomal spatial positioning and the regulation of myogenic genes in the tumor cells.

## Results

### Rhabdomyosarcoma cells exhibit similar patterns of nuclear shape and size characteristics as human myoblasts

To assess differences in nuclear characteristics between normal human myogenic cells and rhabdomyosarcoma (RMS) cells, cultured proliferating human myoblasts (MB), differentiated myotubes (MT) and cell culture models of RMS representing both PAX-FOXO fusion-negative (RD, SMS-CTR) and fusion-positive (RH30) subtypes had their nuclei visualized using DAPI staining (**Fig 1A**). Given all cell lines were grown in adherent conditions and consistently exhibited relatively little depth (typically ∼3 - 4 μm, data not shown) their characteristics were considered as projections into only the x- and y-axes, rather than as three-dimensional structures.

**Figure 1.**
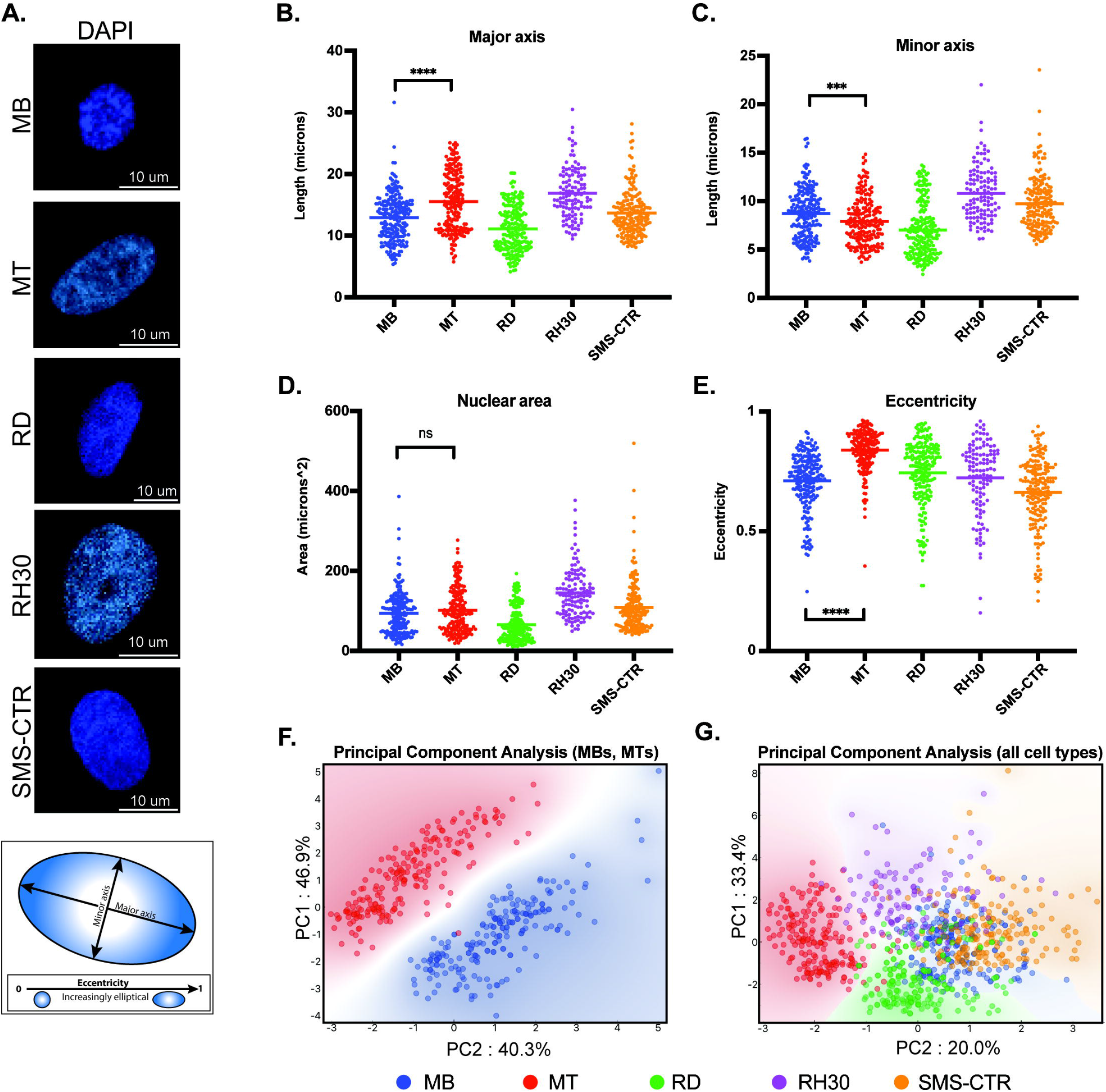
Nuclear shape and size of rhabdomyosarcoma cell lines are more similar to myoblasts than myotubes. A) Representative microscopy images of the DAPI-labeled nuclei of primary human myocytes and rhabdomyosarcoma cell lines. Scale bar as indicated. Diagram below shows representative major and minor axes in a nucleus. B) Scatterplot demonstrating the major axis length of each of the indicated cell types. Each point indicates the measurement from a single nucleus, with the mean indicated by the horizontal line. Only the statistical testing between MB and MT cells is indicated. C) Scatterplot demonstrating minor axis length measurements as in B. D) Scatterplot demonstrating area measurements, as for B. E) Eccentricity measurement scatterplot, as for B. F) Principal component analysis was performed using the measurements from B – E for myoblasts (MB) and myotubes (MT) only, and demonstrates a strong separation between the cell types based only on those nuclear characteristics. Each axis label (PC1 and PC2) indicates the respective percentage of variability explained. G) When the three rhabdomyosarcoma cell lines are included in the PCA, overlap is seen between the myoblasts (MB) and rhabdomyosarcoma cells, but not the myotubes (MT) and rhabdomyosarcoma cells. As for 1F, the respective percentage of variability that is explained is listed on each axis. Statistical testing done by t-tests with unequal variance. ns: not significant; ^***^: p <0.001; ^****^: p <0.0001, n = 212 (MB), 199 (MT), 214 (RD), 121 (RH30), 181 (SMS) nuclei.

Nuclear characteristics including length of major and minor axes (**Fig 1B, 1C**), nuclear area (**Fig 1D**) and eccentricity (**Fig 1E**) were determined for each cell type. Differentiation of myoblasts into myotubes resulted in an average increase of major axis length from 12.8 to 15.5 μm (**Fig 1B**), an average decrease of minor axis length from 8.7 to 7.9 μm (**Fig 1C**), no difference in nuclear area (**Fig 1D**), and an average increase in eccentricity from 0.71 to 0.84 (**Fig 1E**). Principal component analysis (PCA) of the normal myogenic cells demonstrated that these basic nuclear characteristics were sufficient to distinguish proliferating myoblasts from differentiated myotubes with a high degree of fidelity and account for 87% of the variability in the data (**Fig 1F**).

While RMS cell lines typically exhibited similar distributions as the normal cells for their nuclear characteristics, as demonstrated by comparable coefficients of variation for individual characteristics, (**Suppl Table S1**) their average values frequently differed both between tumor cell lines (**Fig 1B and C**, consider RD versus RH30) and when tumor cells were compared to the normal cells (**Fig 1C**, consider RD relative to MB and MT). The characteristic that all RMS cell lines demonstrated a fairly consistent pattern with was eccentricity, where they universally exhibited lower eccentricity compared to human myotubes (**Fig 1E**). PCA analysis that included both RMS cells and the normal myogenic cells demonstrated that, while the variability that was accounted for decreased to 53%, all RMS cell lines showed overlap with the proliferating MBs, while differentiated MTs demonstrated separate clustering (**Fig 1G**).

### Chromosomes 2 and 18 exhibit distinct patterns of nuclear area occupancy and radial localization in myogenic and RMS cells

A subset of human chromosomes have previously been identified as having a greater number of myogenic tandem gene arrays (TGAs) - comprised of linear stretches of genes differentially regulated during the process of myogenesis^9^ – than would be expected by chance. We chose to interrogate the differences in a variety of chromosome organizational characteristics in normal myogenic and RMS cells in a chromosome with a significant enrichment for myogenic TGAs (Chromosome 2) and a chromosome without such enrichment (Chromosome 18). For both chromosomes, the same cell lines examined in Figure 1 were hybridized with chromosome paints to allow their visualization alongside DAPI staining to delineate their nuclei (**Fig 2A and C**). In all cell lines, the chromosomes typically occupied all or the majority of the z-axis (**Suppl Fig S1**) so, as for Figure 1, imaging was projected into two dimensions prior to analysis. Imaging analysis universally identified two chromosome signals per nucleus for the myogenic cells, while variable numbers were identified in tumor cells (**Figs 2A and C and Suppl Fig S2A and B**), consistent with chromosomal fragmentation and/or aneuploidy.

**Figure 2.**
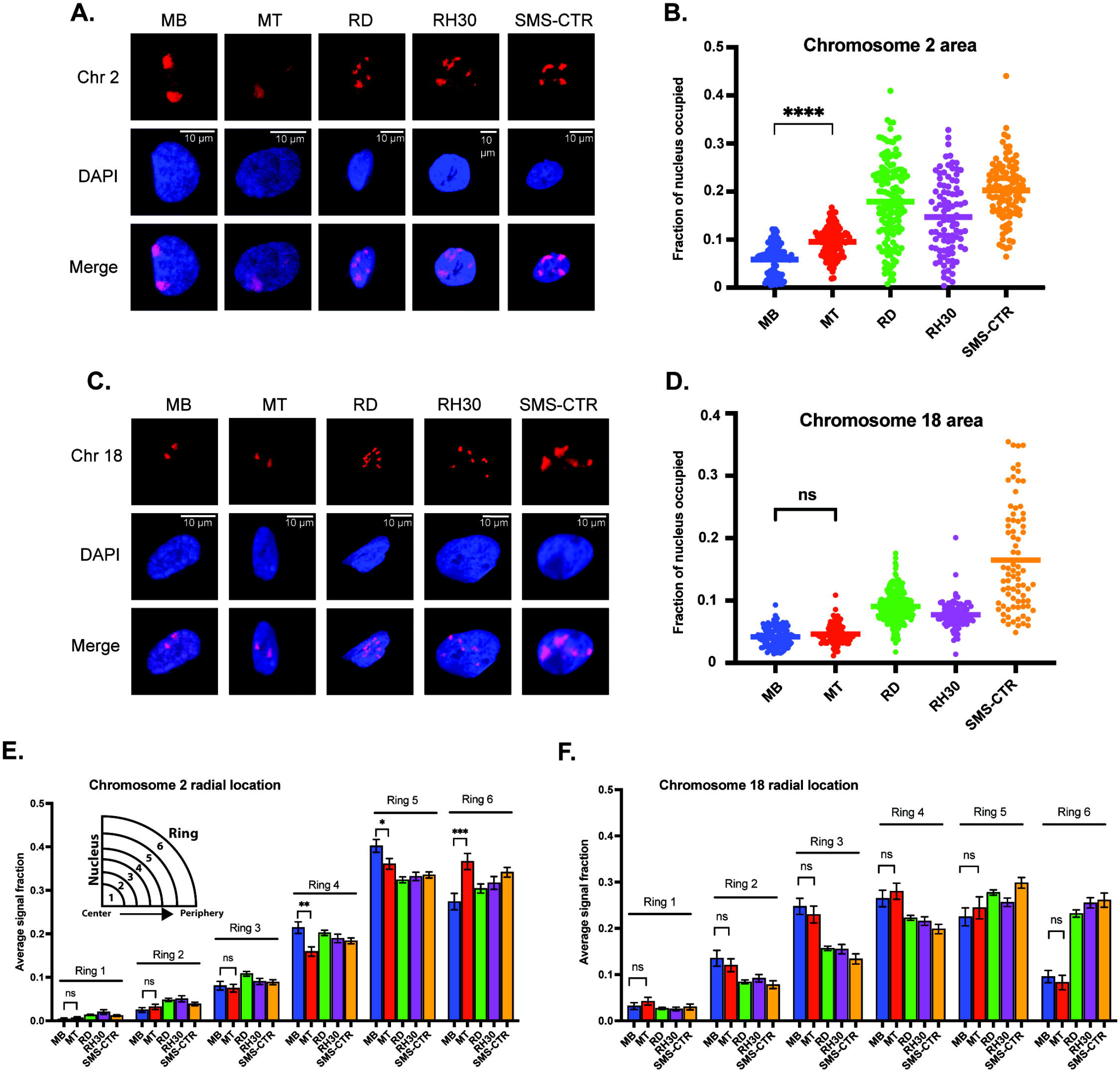
Chromosome-specific differences in organizational characteristics with differentiation and in tumor cells. A) Representative microscope images of myoblasts (MB), myotubes (MT) and rhabdomyosarcoma cell lines with chromosome 2 visualized through the hybridization of a chromosome paint. Images represent visualization of either the paint fluorophore (Chr 2), the nucleus (DAPI), or the merge of the above, as indicated to the side. B) The area of the nucleus occupied by chromosome 2 was determined for each cell type, and is represented as a scatterplot, with each nucleus measured represented as a single point, and the mean represented by the horizontal bar. Statistical testing is shown only for the MB to MT comparison and demonstrates a significant increase in area occupied by chromosome 2 in the differentiated cells. C) Representative microscope images for chromosome 18 visualization as in 2A. D) Plots of the area of the nucleus occupied for chromosome 18, as in 2B. Statistical testing of the means of MBs and MTs demonstrated no significant difference in area occupied.E) Graphing of the average fraction and SEM of chromosome 2 fluorescent signal present in each one of six concentric nuclear rings (ring 1: most central, ring 6: most peripheral) demonstrates that in all cell types analyzed, chromosome 2 had little presence in the inner rings compared to the outer rings, and that there was a shift to increased signal presence in the outermost ring with differentiation from MB to MT. F) Graphing, as in 2E, of the average fraction of chromosome 18 fluorescent signal in each of six concentric nuclear rings demonstrates more central localization compared to chromosome 2, and no difference in radial localization with the process of differentiation. ns: not significant; ^*^: p <0.05; ^**^: p <0.01; ^***^: p <0.001; ^* ** *^: p <0.0001. n = 100 – 143 nuclei per cell type for chromosome 2; 78 – 270 nuclei for chromosome 18.

The area of the nucleus occupied by chromosome signal showed differences both in normal myogenic cells, as well as between normal and tumor cells. While nuclear area occupancy of chromosome 2 nearly doubled on average as myogenic cells differentiated (MB: 5.9%, MT: 9.5% of nuclear area occupied), the area occupied by chromosome 2 in the tumor cells was variable from cell to cell, but with notably higher averages than the normal cells (**Fig 2B**). In contrast, chromosome 18 showed no difference in nuclear area occupied during normal myogenesis (**Fig 2D**, compare MB to MT), and while all tumor cell lines again demonstrated higher averages, for 2 of the 3 lines, the clustering around the mean was notably more consistent than for the last cell line (**Fig 2D**, compare RD/RH30 to SMS-CTR). For each chromosome, area occupied relative to the extent of chromosomal fragmentation was plotted, and showed no evidence of a consistent relationship (**Suppl Fig S2A and B**), suggesting that the difference seen in the tumor cells is not a function of fragmentation and/or aneuploidy.

In addition to the area occupied, the radial positioning of each chromosome was determined as a function of the chromosome signal seen in each of six concentric rings per nucleus. Chromosome 2 exhibited very low average presence in the innermost nuclear rings (rings 1 and 2), moderate presence in the middle rings (3 and 4) and the most presence in the outermost rings (rings 5 and 6) (**Fig 2E**). Myogenic differentiation resulted in chromosome signal changes in rings 4-6, with a decrease in signal in both rings 4 and 5 (from 0.21 to 0.16 and 0.40 to 0.36, respectively), with a concomitant increase in signal in ring 6 (from 0.27 to 0.37), consistent with chromosome 2 shifting to a more peripheral localization with differentiation (**Fig 2E**, compare MB and MT bars). In contrast, while chromosome 18 also demonstrated extremely low occupancy in ring 1, it was seen more frequently in ring 2 and both the middle rings compared to chromosome 2, with no change in response to differentiation (**Fig 2F**).

When considering the radial localization in the tumor cell lines, notable differences were seen between the two chromosomes. Chromosome 2 localization did not clearly align more consistently with either myoblasts or myotubes (**Fig 2E**, consider ring 5, where all tumor lines are lower than both myogenic cell types and ring 4, where all tumor lines are instead intermediate between the myogenic cells), but all tumor cell lines demonstrated localization that was relatively comparable to that seen in the normal cells (**Fig 2E**). In contrast, chromosome 18 often exhibited substantial discrepancy between normal and tumor cells. Chromosome 18 signal was notably lower in rings 2-4 in all tumor cells compared to normal, and much higher in ring 6 compared to the normal cells (**Fig 2F**), consistent with a shift to more peripheral localization in the tumor cells. When the radial localization of each chromosome was visualized on a per-cell basis, rather than the cell-type averaging as above, despite cell-to-cell variability for each cell type, the same overall patterns as described above were clearly visible (**Suppl Fig S3A and B**).

In summary, in normal myogenesis, the chromosome enriched for myogenic TGAs, chromosome 2, exhibits an increase in the nuclear area it occupies and a shift to a more peripheral localization with differentiation, while the non-enriched chromosome 18 changes neither its area nor its peripheral localization. In tumor cell lines, both chromosomes exhibit an increase in area relative to normal cells, but while chromosome 2 appears to maintain a fairly preserved radial occupancy compared to the normal cells, chromosome 18 has a substantial change in its positioning in all tumor lines tested when compared to normal.

### Nuclear density and chromosomal positioning

Given the changes seen in radial positioning in normal cells for chromosome 2, and between normal and tumor cells for chromosome 18, we sought to determine if there was a detectable change in the positioning of either chromosome relative to areas of differing DNA density that could potentially correspond to large-scale regions of hetero-or euchromatin. Using DAPI intensity as a marker for DNA density, we sought to determine both whether large-scale variations in DNA density were observed, and whether there were differences in the overlap between chromosomal signal and DNA density between normal and tumor cells.

The average DAPI intensity was first calculated for each cell type as a function of the same six concentric nuclear rings used to determine radial chromosome localization. Regardless of the cell type analyzed, increasing DAPI intensity was consistently observed as concentric rings became more peripheral (**Fig 3A**). To account for the possibility that the tumor cells may possess significant cell-to-cell variability, but relatively consistent averages when compared to normal cells, the relative DAPI intensity in each ring was plotted for each individual nucleus and demonstrated a pattern consistent with that seen for the average value analysis, with increases in intensity across the innermost and middle rings, and highest intensity in rings 5 and 6, relatively comparable between the two (**Fig 3B**).

**Figure 3.**
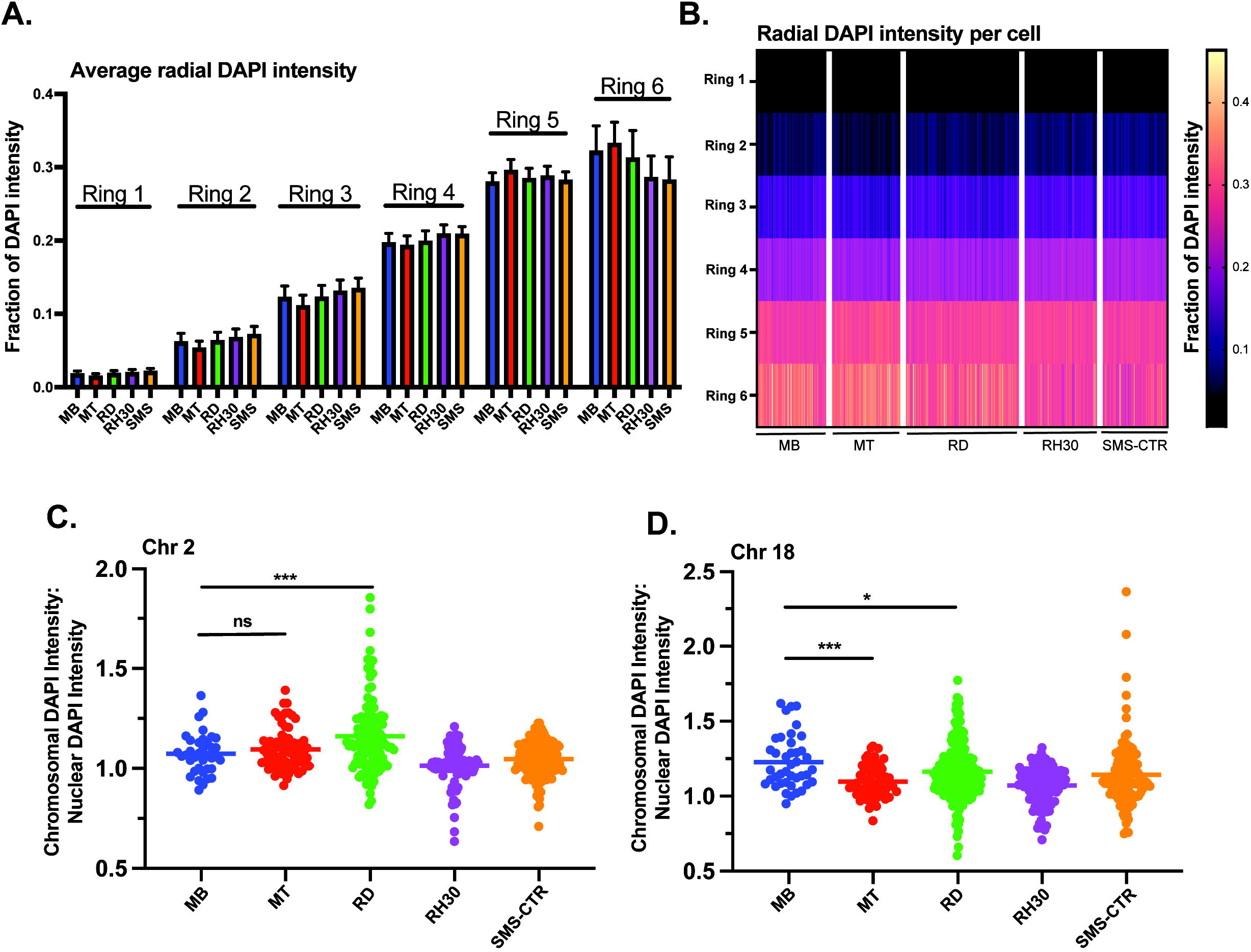
Myogenic and RMS cells exhibit similar overall DNA density patterning, but cell- and chromosome-specific differences in chromosome DNA density. A) DAPI intensity in nuclei as a function of radial distance is shown divided up across six concentric nuclear rings, where ring 1 is the innermost portion of the nucleus and ring 6 the most peripheral. Each bar represents the average intensity for a given cell type in a given ring, with error bars indicating the standard deviation. B) The data from 3A shown here as a heat map, with each column representing a single individual nucleus. C) A scatterplot depicting the ratio of the DAPI intensity of the region of the nucleus occupied by chromosome 2 relative to the normalized DAPI intensity of the whole nucleus for each cell type as indicated. Each point represents a single chromosome, the mean is indicated by the horizontal line. D) A scatterplot depicting chromosome 18-specific DAPI intensity relative to normalized nuclear DAPI intensity as in 3C. Statistical testing done by t-tests with unequal variance. ns: not significant; ^*^: p <0.05; ^***^: p <0.001; n = 96 – 167 nuclei per cell type for A and B; n = 37 – 167 chromosomes per cell type for chromosome 2; 41 – 228 chromosomes for chromosome 18 for C and D.

To investigate the relationship between chromosomal positioning relative to DNA density, the ratio of the DAPI intensity of the region of the nucleus occupied by a given chromosome to the normalized DAPI intensity of the nucleus as a whole was determined and plotted for both chromosomes 2 and 18 (**Fig 3C and D**). Chromosome 2 exhibited no significant difference comparing MBs to MTs (means of 1.07 and 1.1, respectively) while the RMS lines exhibited variable ratios, with RDs exhibited higher average values compared to MBs (mean of 1.16), consistent with chromosome 2 in those cells occupying areas of greater DAPI intensity, RH30s exhibiting lower average values compared to MBs (mean of 1.01) and SMS-CTRs were statistically indistinguishable from MBs (mean of 1.05). Chromosome 18, in contrast, did exhibit a difference between MBs and MTs, with chromosome 18 in MTs occupying areas of lower DAPI intensity on average, as did all tumor cell lines (**Fig 3D**).

### Myogenic cells exhibit differentiation-dependent preferential positioning of chromosome 2 along the major nuclear axis that is absent in rhabdomyosarcoma cells

Since myogenesis affects the inter-allelic distance of the two alleles of the myogenic transcription factor myogenin^9^, we investigated whether there were differentiation dependent differences in inter-chromosomal distances and angles (**Fig 4A**). While chromosomes 2 and 18 exhibited chromosome-specific patterns in average inter-chromosomal distances, with both chromosomes 18 frequently being located more closely together than chromosomes 2, for a given chromosome, no difference in average distance was seen between myoblasts and myotubes (**Fig 4B**). Similarly, no difference was noted in the inter-chromosomal angles with differentiation (**Fig 4C**), suggesting that there is no coordinated regulation of the positioning of either chromosome relative to each other with differentiation.

**Figure 4.**
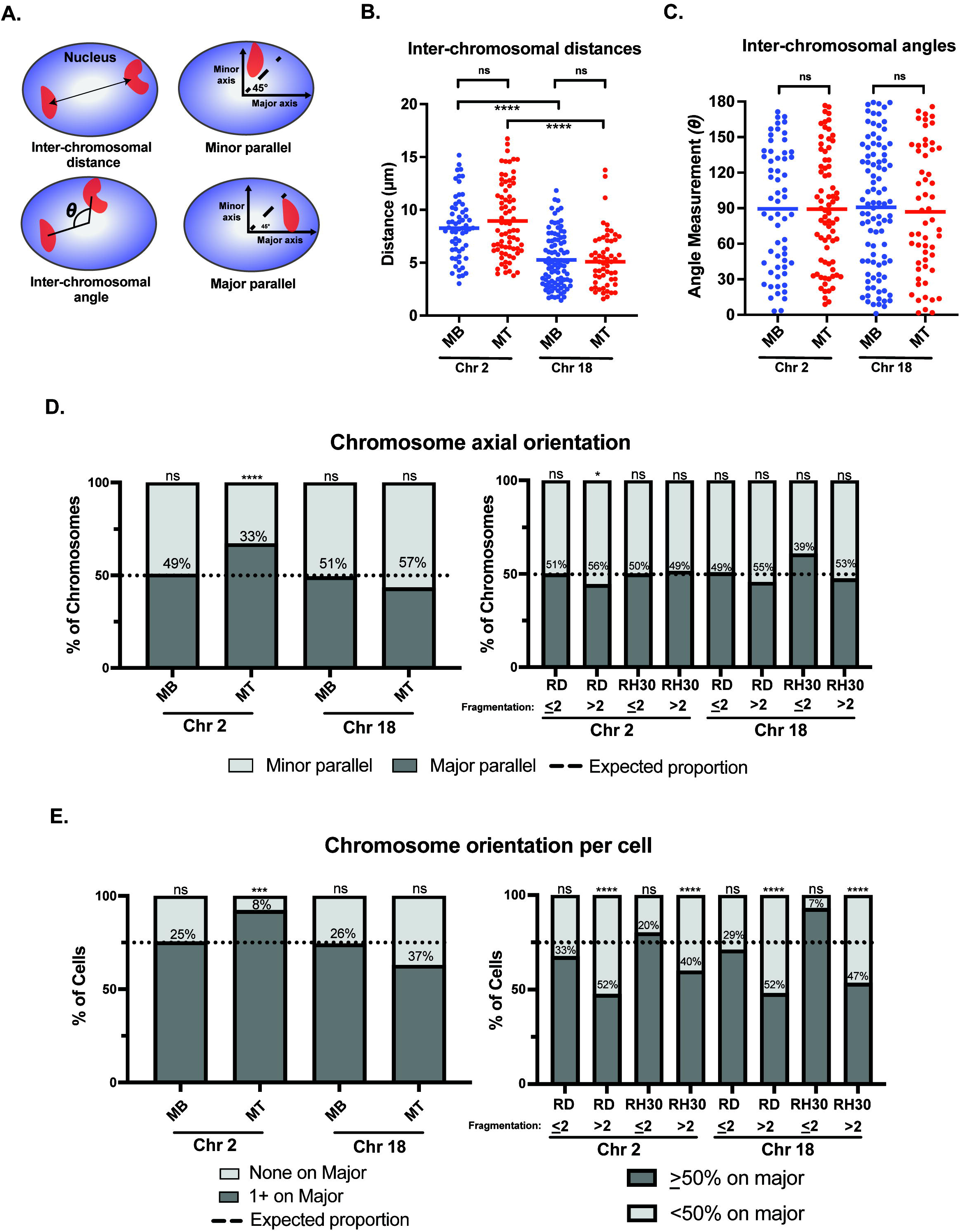
A chromosome-specific change in positioning relative to the nuclear axis occurs with normal myogenic differentiation and is absent in rhabdomyosarcoma cells. Diagrammatic depictions of inter-chromosomal distance, inter-chromosomal angle, and minor and major parallel of chromosomal signal. B) A scatterplot representing the inter-chromosomal distance between the centroids of either chromosome 2 or chromosome 18 (indicated below) in the nuclei of either myoblasts (MB) or myotubes (MT). Each point represents the measurement in a single nucleus, and the horizontal line represents the mean. C) The scatterplot shows the inter-chromosomal angle for the same nuclei and chromosomes as measured in 4B. D) Bar graphs that indicate the percentage of all measured chromosomes for an indicated cell type that were assigned either as being associated with the minor axis (minor parallel) or the major axis (major parallel), as depicted in 4A. The dotted line indicates the expected proportion of chromosomes that would be found associated with a given axis if they were organized in space at random. E) Bar graphs for the same cells and measurements as in 4D, but with the calculation done as the orientation of the chromosomes in each individual nucleus. Nuclei in normal cells were classified either as having no chromosomes assigned to the major axis (none on major) or at least 1 chromosome assigned to the major axis (1+ on major). In tumor cells, cells were grouped as either having <50% of their identified chromosomal signals on the major axis, or >50% of chromosomal signals on the major axis. As in 4D, the dotted line indicates where the expected division would be between the two classifications if each chromosome was randomly positioned in nuclear space. In both D and E, the number within the bars indicates the value assigned to the minor parallel for each condition. Statistical testing in B and C was by t-tests with unequal variance; D and E was by using a binomial test to compare observed count frequencies. ns: not significant; ^*^: p <0.05; ^***^: p <0.001; ^** **^: p <0.0001; n = 55-93 nuclei per cell type : chromosome pairing in B and C; n = 80 – 140 nuclei per cell type for chromosome 2 in D and E; n = 60 – 220 nuclei per cell type for chromosome 18 in D and E.

In contrast however, a chromosome-specific, differentiation-dependent difference was seen when assessing chromosomal positioning relative to the nuclear axis. When considering where chromosomes were located relative to the major and minor axis of the nucleus (**Fig 4A**), chromosome 2 in myoblasts showed positioning that was indistinguishable from what would be expected by chance (**Fig 4D, leftmost bar**), even if taking into account corrections for the average greater length of the major axis relative to the minor axis in those cells (**Suppl Table S2**). In contrast, in myotubes, chromosome 2 was significantly more frequently identified along the major axis than would be expected to occur by chance, despite the even greater average major axis length in those cells compared to myoblasts (**Fig 4D**, consider Chr 2 MT bar). This relationship was also seen when chromosome positioning relative to the nuclear axis was considered on a per-cell basis (**Fig 4E, left panel**), with <10% of myotube nuclei having both chromosomes 2 located along the minor axis, less than half the percentage that would be expected by chance if chromosomes were randomly and independently distributed in nuclear space. This relationship continued to be statistically significant, even when accounting for axial length and area discrepancies (**Suppl Table S2**). Chromosome 18, on the other hand, whether analyzed on a per-cell basis or for all the chromosomes as a whole, was found at proportions that were statistically indistinguishable from what would be expected by chance in both myoblasts and myotubes (**Fig 4D and E, left panels**).

Given the complexity of computing expected probabilities for tumor cells with fragmented chromosomes, the tumor nuclei were dichotomized into two groups for the axial analysis: those cells with two (or fewer) detectable chromosomal signals per nucleus and those with greater than 2 detectable chromosomal signals. Analysis of the distribution on the per chromosome basis showed only one condition that was significantly different from the distribution that would be expected by chance, with high fragmentation score RD cells having a slightly increased rate of chromosome 2 being aligned along the minor parallel (**Fig 4D, right panel**). When considering the per-cell distributions, results were consistently related to the fragmentation score: all chromosome-cell type pairs with a fragmentation score of <2 demonstrated a distribution indistinguishable from the expected proportion, while all chromosome-cell type pairs with a fragmentation score of >2 were significantly different from that proportion (**Fig 4E, right panel**). Overall, the distribution of positioning in high fragmentation cells demonstrated a relatively even distribution between major and minor axis positioning (**Fig 4E, right panel**, consider RD >2 Chr 2 and RD >2 Chr 18).

### Induction of differentiation in RD cells increases positioning of Chromosome 2 along the major nuclear axis

With the RMS cell lines exhibiting patterns of chromosome 2 axial positioning similar to what was seen in myoblasts, as opposed to the preferential positioning seen in myotubes, we tested whether we could restore chromosome 2 to major axis positioning via the induction of differentiation in RMS cells. The introduction of the pro-myogenic miRNA miR-206 has previously been shown to lead to differentiation and cell cycle withdrawal in RMS cells^14-16^. RD cells were transiently transfected (**Suppl Fig 4A**) with either a miRNA mimetic of miR-206 or a negative control mimetic. As expected, incubation of transfected cells in low-serum differentiation media led to morphologic change and significant expression of the marker of myogenesis, myosin heavy chain (MHC), in cells transfected with the miR-206 mimetic but not the negative control mimetic (**Fig 5A**). Visualization of chromosome 2 in transfected cells worked as it had for untreated cells (**Fig 5B**), and the axial orientation of chromosomes was assessed on both a per chromosome (**Fig 5C**) and per cell (**Fig 5D**) basis. In both analyses, a statistically significant increase in the proportion of chromosomes along the major axis was found overall in those cells transfected with the miR-206 mimetic compared to those transfected with the negative control mimetic, with the per chromosome proportion increasing from 45% to 55% and the per cell proportion increasing from 48% to 75% (p = 0.0089 and 0.0024, respectively). Analysis based on fragmentation status was performed and showed no clear difference in proportions, though it was limited due to a small number of cells that met criteria for low fragmentation status, especially in the miR-206 mimetic condition (data not shown).

**Figure 5.**
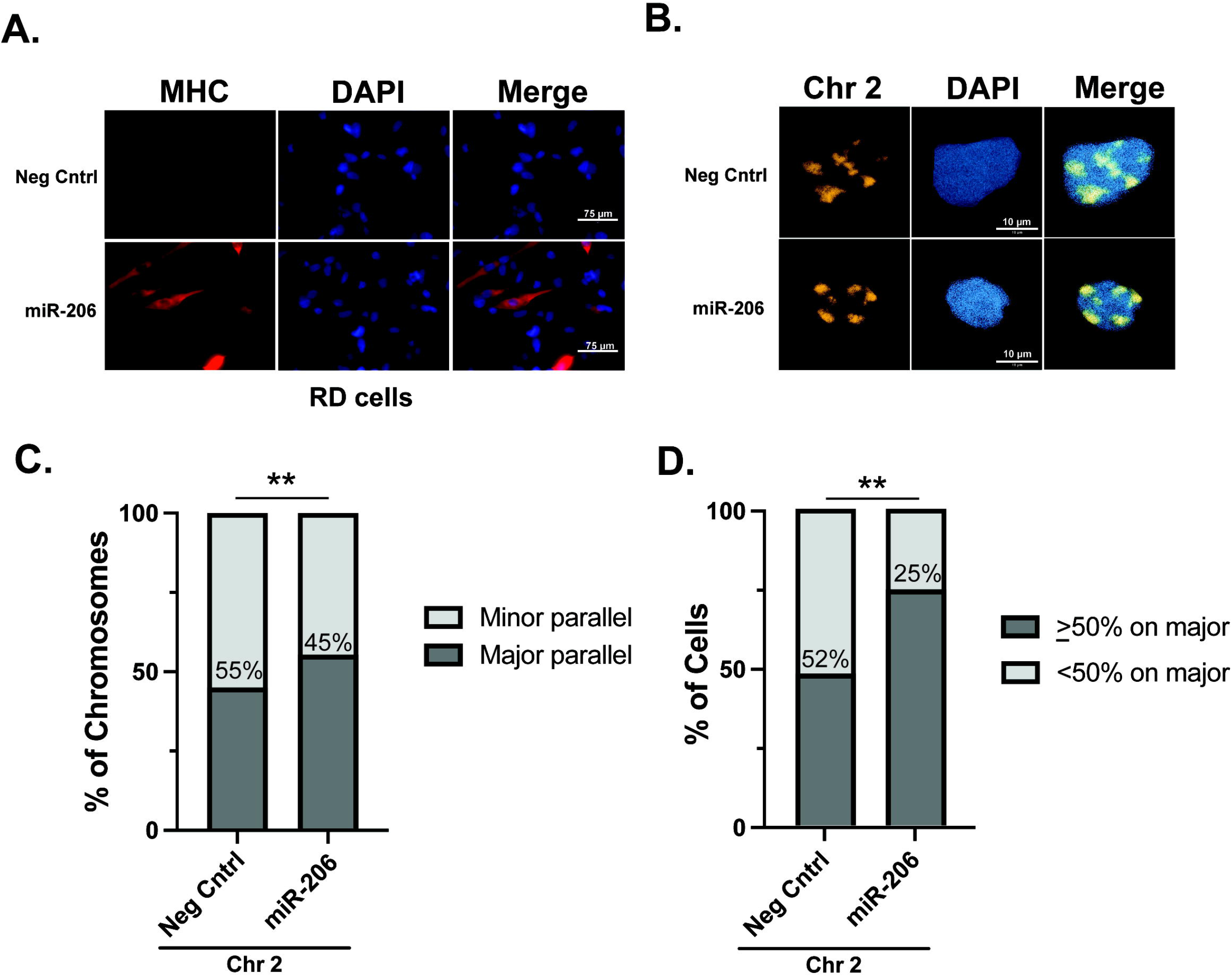
Differentiation of RD cells via transfection of a miRNA miR-206 mimetic increases the frequency of chromosome 2 positioning along the major nuclear axis. A) Immunohistochemistry for myosin heavy chain, a marker of myogenesis, shows an increase in its expression and the formation of elongated cellular shapes, some of which exhibit multi-nucleation, both consistent with differentiation of RMS cells specifically in those cells transfected with the miR-206 mimetic and not the negative control mimetic. B) Representative images of chromosome 2 visualization via hybridization of chromosome paint in RD cells. C) The proportion of chromosome 2 assigned as being associated either with the minor axis (minor parallel) or the major axis (major parallel) in RD cells transfected as indicated. The number within the bars indicates the value assigned to the minor parallel for each condition. D) The proportion of cells with chromosome 2 positioning as indicated on a per-cell basis. The percentage of cells with <50% of chromosome signals along the major axis is indicated within the bars. For both 5C and 5D, statistical testing was performed using Fisher’s exact test for proportions. ^* *^: p<0.01. n = 76 and 61 nuclei for negative control and miR-206 nuclei, respectively.

The nuclear characteristics of the transfected cells were assessed, to determine if miR-206 treatment would result in nuclei taking on myotube nuclei characteristics. There was no evidence of difference in the nuclear minor axis length, but major axis, area, and eccentricity all decreased on average with differentiation. For both of the axial length measurements and for the area, miR-206 transfection led to more homogeneity between nuclei, as represented by reduced coefficients of variation (**Suppl Fig S4B-E**). There was a small but statistically significant shift to more internal localization of chromosome 2 in the miR-206 transfected cells, with a 2% decrease in the average signal in the outermost ring (6), and increases of only approximately 1% in rings 1-3 (**Suppl Fig 4F**).

## Discussion

Previous work examining the role of specific chromosomes in myogenic cells has demonstrated that a subset of chromosomes are enriched for tandem gene arrays (TGAs) of genes that are differentially regulated during normal myogenesis. We now demonstrate that differential behavior in chromosome organization and spatial positioning occurs between a chromosome enriched for such TGAs (Chromosome 2) and one that is not enriched for them (Chromosome 18) during normal myogenesis. Most strikingly, we identify a preferential spatial positioning of chromosome 2 along the major nuclear axis in differentiated myotubes that is absent in both proliferating myoblasts, and baseline RMS cells, but can be partially restored in RMS cells that are induced to differentiate.

To obtain a basis for analyzing chromosomal topologies between normal myocytes and RMS cells, we characterized their nuclei on a variety of features, including axial measurements, area, and eccentricity. Others have demonstrated changes in myocyte nuclear volume and thickness with normal myogenesis^28 29^, with myotubes more ‘flattened’ and elongated with differentiation. While our analysis in two dimensions demonstrates no change in nuclear area, myotubes exhibit an increase in one dimension accompanied by a concomitant decrease in the other and an increased eccentricity, consistent with the previously described changes. The RMS cells exhibit notable variability both in the absolute values of their nuclear characteristics, and their relative relationship to the normal cell measurements. For all three RMS cell lines, their eccentricity was closer to myoblasts than myotubes, demonstrating a more rounded shape. Principal component analysis demonstrated that those four nuclear characteristics alone allowed for excellent discrimination of the undifferentiated from differentiated myocytes, and clustered the RMS cells with myoblasts, and distinctly apart from myotubes.

Interestingly, our induced differentiation of the RMS cells with the microRNA miR-206 led to changes in nuclear characteristics, but rather than the RD cells becoming more elliptical and elongated with differentiation as the normal myocytes did, they instead became smaller and more rounded. It is unclear if that is because 1) despite differentiation, the RMS cells do not (or cannot) appropriately regulate their nuclear morphology and size, 2) the changes in nuclear characteristics would ultimately occur if the RMS cells were kept in differentiation-promoting conditions for long enough, or 3) the fact that the transfected RMS cells were cultured on poly-Llysine coated surfaces to tolerate subsequent hybridization steps without excessive cell loss. The poly-L-lysine did result in some alteration of nuclear size characteristics at baseline, as control transfected RD cells possessed somewhat difference mean nuclear axes measurements compared to non-transfected RD cells. Notably, while the range of eccentricity measurements was equivalent between control and miR-206 transfected cells, all three of the nuclear size characteristics became more homogeneous with induced differentiation. While the presence of the microRNA itself could not be directly tracked in individual cells, the increased homogeneity of the entire population of nuclei in the miR-206 condition agrees with the high efficiency of transfection of RD cells we saw as measured by transfection of a fluorescently labeled oligo. This in turn suggests that the expression of markers of myogenesis, such as myosin heavy chain, are separate - either temporally or in regards to regulation – from the processes that reshape the cell nucleus, as myosin heavy chain expression was not seen as widely in the miR-206 transfected cells as the changes in nuclear morphology were.

Chromosomal positioning in the nucleus has been shown to be cell-type specific^30 31^, and the positioning of centromeres has been shown to change with myogenesis^10^, with a subset of centromeres exhibiting differential radial positioning, including chromosome 2^29^. Using labeling of the entire chromosome, our data also identify differential radial positioning with myogenesis for chromosome 2, accompanied by a near doubling in the area of the nucleus the chromosome occupies. While the increased nuclear area of occupancy of chromosome 2 seen with differentiation is potentially consistent with this myogenic TGA-enriched chromosome taking on a more ‘open’ chromatin conformation to permit increased transcription of myogenic genes, we could find no evidence of the chromosome as a whole occupying an area of the nucleus with lower DNA density with differentiation. Possibilities for this finding include 1) that use of DAPI as a marker of DNA density is insufficiently sensitive to see a difference that is actually present and labeling of specific epigenetic marks (such as H3K4me3) could reveal differences, 2) that changes are present but localized to one or more portions of the chromosome and treating the chromosome as a single region for the analysis masked the difference in those areas, and/or 3) that the shift of the chromosome to a more peripheral localization, where we consistently saw higher DNA density in the DAPI staining, confounded the analysis similarly to possibility #2. More sensitive techniques, such as the use of sequencing techniques to assess DNA accessibility and contacts, could help delineate between the possibilities and relate them to regulation of the myogenic genes located on the chromosome.

The chromosome-specific differences seen in peripheral localization between the normal and RMS cells is particularly notable, especially when considered alongside the area of occupancy differences. For both chromosomes 2 and 18, RMS cells universally occupied a larger nuclear area compared to normal cells, but exhibited notably disparate behaviors in regards to localization. While chromosome 2 radial positioning was quite similar between normal and RMS cells, chromosome 18 exhibited large differences, particularly notable when examining the occupancy of the most peripheral ring, ring six. Given the association between nuclear positioning and gene regulation^7 32-34^, it’s tempting to speculate that the enrichment for myogenic TGAs on chromosome 2 results in RMS cells continuing to appropriately regulate the positioning of that chromosome, and appropriate regulation of chromosome 18, with its lack of enrichment for myogenic TGAs, being lost. If so, we would expect it to be a more general pattern that would be seen when the radial positioning of other chromosomes is determined in both normal and RMS cells. It also suggests that analysis of gene regulation and expression in RMS cells considered on the basis of the spatial positioning of the chromosomes they reside on, rather than by clustering genes on the basis of similar function or biological processes, would demonstrate chromosome-specific differences.

Our identification of differentiation-dependent preferential positioning of chromosome 2 along the major nuclear axis raises the possibility that a specific subset of chromosomes experience unique stress during myogenesis. Changes in nuclear shape – such as occurs during myogenesis – have been associated with increases in DNA damage and are related to actin activity^35^. Increased DNA damage, in turn, is associated with the generation of gene-gene fusions^36^. The increased frequency of chromosome 2 positioning in a specific nuclear spatial location may subject it to a different set of forces compared to a chromosome without such preferential positioning. Given that chromosome 2 is one of the chromosomes that can be involved in the characteristic PAX-FOXO gene fusions seen in a subset of RMS tumors, it suggests the possibility that the spatial positioning of chromosome 2 may contribute in some manner mechanistically to the formation of the PAX-FOXO fusions. Regardless of that possibility, the increased alignment of chromosome 2 along the major nuclear axis with induced differentiation of the RMS cells suggests a relationship between chromosomal spatial positioning and the regulation of myogenic genes in those tumor cells, though further studies will be necessary to determine if there is a direct causal relationship.

In summary, we have identified a differentiation-dependent preferential spatial positioning of chromosome 2 in normal myocytes that is absent in both fusion-negative and fusion-positive RMS cells that is at least partially restored with induced differentiation of the tumor cells. When considered along with our data demonstrating chromosome-specific differences in other chromosomal organizational characteristics such as radial positioning, and the area of the nucleus occupied, it highlights the relevance of considering chromosomal topologies when investigating gene regulation in both normal and tumor cells.

## Methods & Materials

### Cell Culture

Cell lines were obtained from ATCC and experiments were performed in cells at low passage number. RD cells were cultured and maintained in DMEM media (Gibco) supplemented with 10% fetal bovine serum and 1% PS (penicillin-streptomycin) (Gibco). RH30 cells were cultured and maintained in RPMI (Gibco) with 10% fetal bovine serum and 1% PS. Primary human myoblasts were cultured and maintained in Mesenchymal stem cell media (ATCC) with the addition of the contents of the primary human skeletal muscle growth kit (ATCC) and 1% PS. Myotubes were generated by having myoblasts reach confluency, and then cultured for 96 hours in differentiation tool media (ATCC) with the addition of 1% PS. For transfected cells, they were plated onto poly-L-lysine coated coverslips which were prepared by submerging glass coverslips in poly-L-lysine solution (MilliporeSigma) for 15 minutes at 37° before rinsing three times with PBS, and then placing under UV light for 15 minutes prior to use.

### Transfections

One day prior to transfection, cells were trypsinized in 0.05% trypsin-EDTA and plated onto poly-L-lysine coated glass coverslips (see above) at sufficient density to reach 50-60% confluency the following day. Cells were transfected using either a miR-206 miRNA mimetic (Thermofisher) or negative control mimetic #1 (Thermofisher) using RNAiMax (Thermofisher). Transfections were performed according to manufacturer’s protocol with the following modifications: the mimetic’s final concentration was 7 nM, no antibiotics were present in the media, and 1.5 uL of RNAiMax was used per well for a 12 well dish. After 24 hours, cells were washed 2x with PBS, and media was changed to low serum differentiation media (DMEM +1% horse serum (Hyclone) + 1x insulin-transferrin-selenium (Corning) + 1% PS). Cells were fixed using 4% paraformaldehyde (PFA) (Electron Microscopy Sciences) in phosphate-buffered saline (PBS) after 48 hours in differentiation media.

For fluorescent oligo transfections, the Block-iT Alexa Fluor Red Fluorescent Oligo (Invitrogen) was transfected at the concentration indicated using RNAiMax as above, with the following modifications. Cells were kept light-protected as much as possible, and 24 hours after transfection, cells were fixed as above, mounted with Diamond antifade with DAPI (Invitrogen), and then visualized using microscopy. Cells were counted manually for positivity for oligo signal and nuclei number.

### Chromosome Paint Hybridization for Adherent Cells

Cells were trypsinized, resuspended in appropriate growth media and then plated onto glass coverslips and allowed to adhere for approximately 16 hours prior to fixation. Cells were rinsed once in PBS prior to fixation in 4% PFA for 10 minutes at room temperature. Cells were then washed 3 times in PBS, followed by incubation in PBS with 0.01% Triton X-100 at room temperature 3 times for 3 minutes, then incubation in 0.5% Triton X-100/1x PBS at room temperature for 15 minutes. Cells were then incubated in 20% glycerol/PBS at room temperature for 3-4 hours. Cells were washed in PBS 3 times for 10 minutes each and then incubated in 0.1 N HCl for 5 min at room temperature. The cells were incubated in 2x SSC twice for 3 minutes each before being placed in 50% formamide (Electron Microscopy Sciences)/2x SSC at room temperature for approximately 18 hours. After the addition of chromosome paints (Metasystems) to the coverslips, slides were heated to 75 degrees C for 2 minutes before being placed at 37 degrees for approximately 72 hours for hybridization. After hybridization, coverslips were washed in 2x SSC washes at 37 degrees three times for 5 minutes each, followed by washes in 0.1x SSC at 60 degrees three times for 5 minutes each. The coverslips were then washed in 4x SSC/0.2% Tween-20 three times for 3 minutes and mounted on microscope slides using Diamond antifade with DAPI (Invitrogen).

### Image Acquisition and Analysis

Images were acquired on a Nikon Confocal Microscope or a Zeiss LSM 800 Confocal microscope. Imaging was done with a Plan Apo VC 100× 1.4 NA oil objective as a multidimensional *z*-stack. The acquired 3D image stacks were then fed through imaging processing pipelines utilizing standard tools on Cell Profiler and Fiji^37 38^. The pipelines performed the functions of translating images to maximal projections, calculating distance, object size, nuclear size and eccentricity, object intensity as a function of nuclear concentric rings, area occupation, object intensity distribution, angle theta, and the centroid of an object.

#### Three-dimensional reconstructions

For 3D reconstruction images, cell images were analyzed with the ZEN blue edition software (version 3.5.093.00008, Carl Zeiss Microscopy GmbH). Z-Stacks were obtained using standard software tools, and 3-D images were rendered by adjusting the brightness within the linear range to allow for optimal chromosome visualization.

#### Axial orientation determination

Using standard Mathematica tools^39^, a script was developed to organize and analyze collected data (https://github.com/luaypurple/Chromosome-Spatial-Analysis). In summary, the orientation of chromosomes along the major and minor axis of their parent cell was assessed by calculating the angles of each chromosome relative to the nuclear center and classifying them based on their orientation. Chromosomal orientation was classified based on the calculated angle between the centroid of the identified chromosomal signal and the determined nuclear axes. Chromosome centroids located between 45° and 135° relative to an axis were grouped separately from those with an angle less than or equal to 45° and greater than or equal to 135°.

Chromosome orientation along the major and minor axis was assessed in two ways: 1) as a summation of all chromosomes analyzed and 2) on a per cell basis. Myoblasts and myotubes were split into two classifications: 1) both chromosomes along the minor axis and 2) one or both chromosomes along the major axis. RMS cells were classified into two groups: 1) greater than or equal to 50% of chromosomes on the major axis, 2) fewer than 50% of chromosomes on the major axis.

#### Nuclear and chromosomal DAPI intensity

To approximate DNA density in chromosome 2 and chromosome 18 regions, we utilized the confocal max-projections obtained as described above with both Chromosome-Paint and DAPI staining. Nuclear borders were defined by DAPI intensity using the in-built Mathematica function FindThreshold utilizing Kapur’s Method of entropy minimization. To account for spatial variations in staining intensity prior to nuclear-mask segmentation, the inbuilt NonlocalMeansFilter function was applied to the DAPI intensity images, normalizing staining over a 1-pixel radius to produce mask boundaries. Composite images with a 2-pixel dilation and 2-pixel erosion were then produced to account for the effect of the DAPI intensity normalization. The resulting masks were then produced and defined as nuclear areas of interest. Areas by this process were excluded if they were greater than 10,000 pixel or if they extended beyond the image boundary. Noting that despite this restriction, multiple adjacent nuclei in close proximity (e.g. multinucleated cells) could still potentially be considered as one nuclear area of interest. Chromosomal masks were produced using Threshold with piecewise garrot partitioning paired with Kapur’s Method, restricting analysis to chromosomes with areas occupying between 100 and 10,000 pixel. To account for Chromosome-Paint staining variation, we utilized a median filter over a 2-pixel radius. As nonbiological effects influence staining intensity (e.g. variability in magnification, illumination intensity, and acquisition time), we restricted our analysis to the ratio of DAPI intensity within a chromosome region of interest compared to the DAPI intensity of the accompanying whole nucleus. Using the inbuilt function, ComponentMeasurements, median DAPI intensity within chromosome regions of interest and accompanying nuclear regions could be obtained. We finally calculated the relative DAPI intensity within the chromosome to the nucleus as follows:

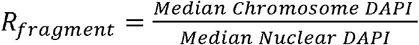

### Statistical Analysis

Statistical testing was performed using Prism 9. Pairwise comparisons were calculated on datasets consisting of, at a minimum, biologically independent duplicate samples using two-tailed t-test with Welch’s correction so as not to assume equal standard deviations in Figures 1 – 4 and Supplemental Figure S4. The regression in Fig S2 was performed using the nonlinear regression tool in Prism 9 and the least squares calculation for the regression. The observed versus expected calculations in Figure 4 used the binomial test in Prism 9 for comparing observed to expected count frequencies. The pairwise comparisons in Figure 5 were performed using Fisher’s exact test in Prism 9. Heat maps were generated using Prism 9’s function. Principal component analysis was performed using Orange3 ^40^.

## Supplemental Figure Legends

**Supplemental Figure S1. Chromosomes 2 and 18 occupy the majority of the z-axis in individual cells**. Representative reconstructions of chromosome signal in the z-axis are shown for chromosomes 2 and 18 in cell types as indicated.

**Supplemental Figure S2. Chromosomal nuclear area occupancy is not a function of chromosome fragmentation**. A) The chromosome fragmentation score (ie. number of detected distinct areas of chromosome paint per nucleus) is shown as a bar and whiskers plot (horizontal line in box: median, ‘+’: mean, whiskers indicate minimum and maximum observed values) for each cell type from the same cells analyzed in Figure 2. As expected, all myoblast (MB) and myotube (MT) nuclei exhibited 2 countable areas of signal per cell, while the tumor cell lines showed variability. Scatterplots below show the fraction of the nucleus occupied per nucleus graphed as a function of the computed fragmentation score for each of the 3 tumor cell lines and a computed regression line. Least squares regression for each cell type showed a negative correlation for RDs, positive correlations for RH30s and SMS-CTRs, and low R2 values overall (as indicated on each graph). B) Chromosome fragmentation scores for chromosome 18 as in S2A, again showing only 2 countable areas of chromosome signal per nucleus for the normal cells, and variable numbers for the tumor cell lines. As for chromosome 2, plots of occupied nuclear area as a function of fragmentation score for each cell type again show significant variability, and very low R2 values with least squares regression. n = 93 – 110 nuclei per cell type.

**Supplemental Figure S3. The pattern of chromosome radial localization in individual cells is similar to what is seen in average measurements**. Heatmaps for the chromosome radial localization of chromosomes 2 and 18 shown for the individual cells that were averaged in Figures 2E and 2F. Each column represents the data from an individual nucleus.

**Supplemental Figure S4. miR-206 transfection in RD cells, nuclear characteristics, and chromosome 2 radial location**. A) RD cells can be transfected at a high rate of efficiency.Transfection of a fluorescently labeled oligo at the concentration indicated results in >80% of cells being scored as positive for signal. Fewer than 5% of control cells (no oligo) are scored as positive using the same criteria. B) The major axis of miR-206 mimetic transfected RD cells are slightly shorter on average compared to cells transfected with a negative control mimetic and have a smaller coefficient of variation (CoV). C) The minor axis of miR-206 transfected cells is not significantly different on average compared to control transfected cells but has a smaller CoV. D) The nuclear area of miR-206 transfected cells is smaller on average compared to control cells, as well as having a smaller CoV. E) miR-206 cells are less eccentric on average, and have a similar CoV to negative control cells. F) Average radial localization of chromosome 2 was computed as a function of six concentric nuclear rings as in Figure 2. All statistical tests were performed as t-tests with unequal variance. ns: not significant; ^*^: p<0.05; ^**^: p<0.01; ^***^: p<0.001; ^****^: p<0.0001; n = 76 and 61 nuclei for negative control and miR-206 nuclei, respectively

## Acknowledgements

KLM is supported, in part, by the National Institutes of Health’s National Center for Advancing Translational Sciences, Grant Number KL2TR001424, as well as the Hyundai Hope on Wheels Scholar Hope Grant, Unravel Pediatric Cancer, the Stanley Manne Children’s Research Institute, and the Ann & Robert H. Lurie Children’s Hospital of Chicago. The content is solely the responsibility of the authors and does not necessarily represent the official views of the National Institutes of Health.

