## Supplementary figures and images for "Differentiation-dependent chromosomal organization changes in normal myogenic cells are absent in rhabdomyosarcoma cells"

### Supplemental Fig S1

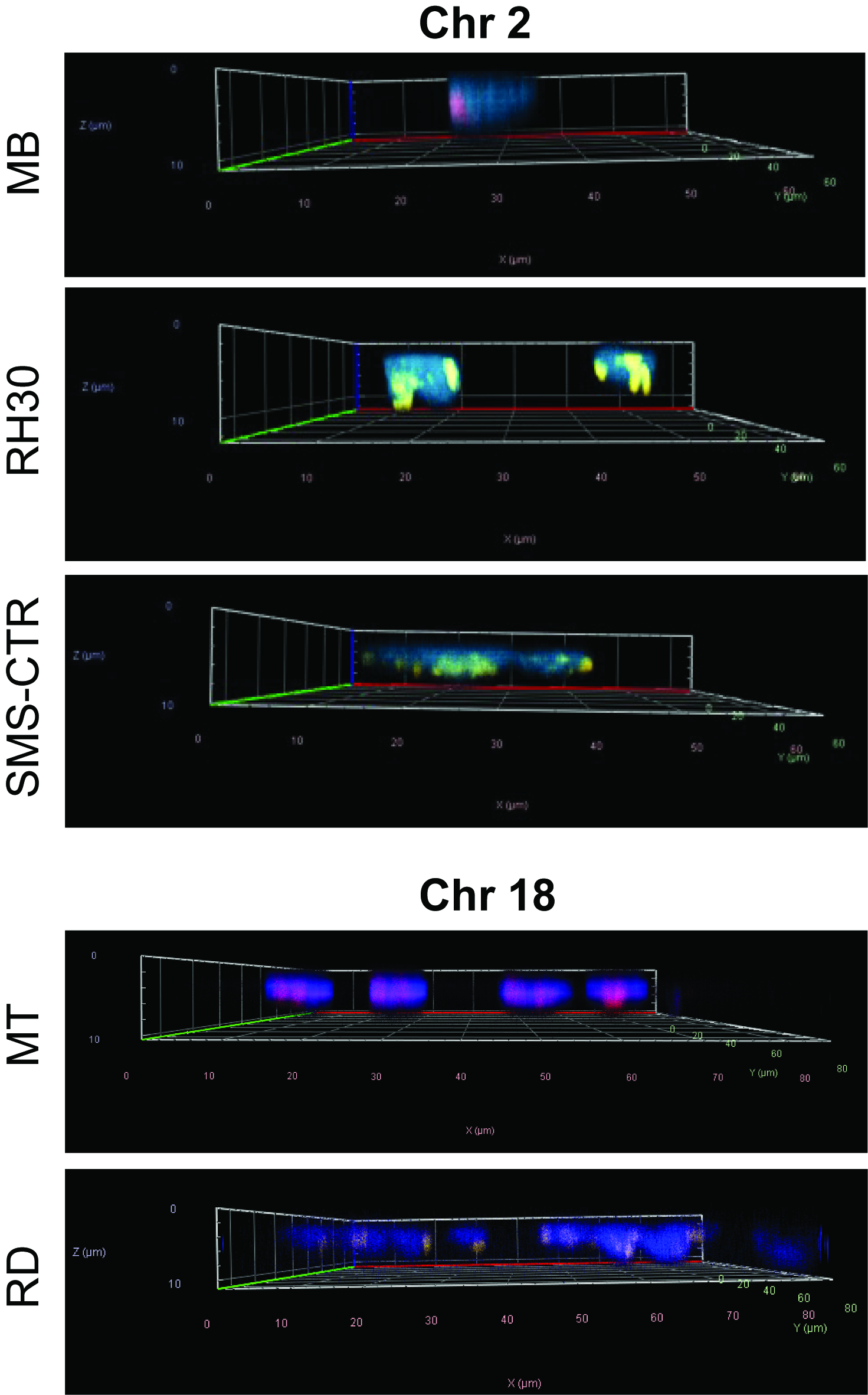

### Supplemental Fig S2

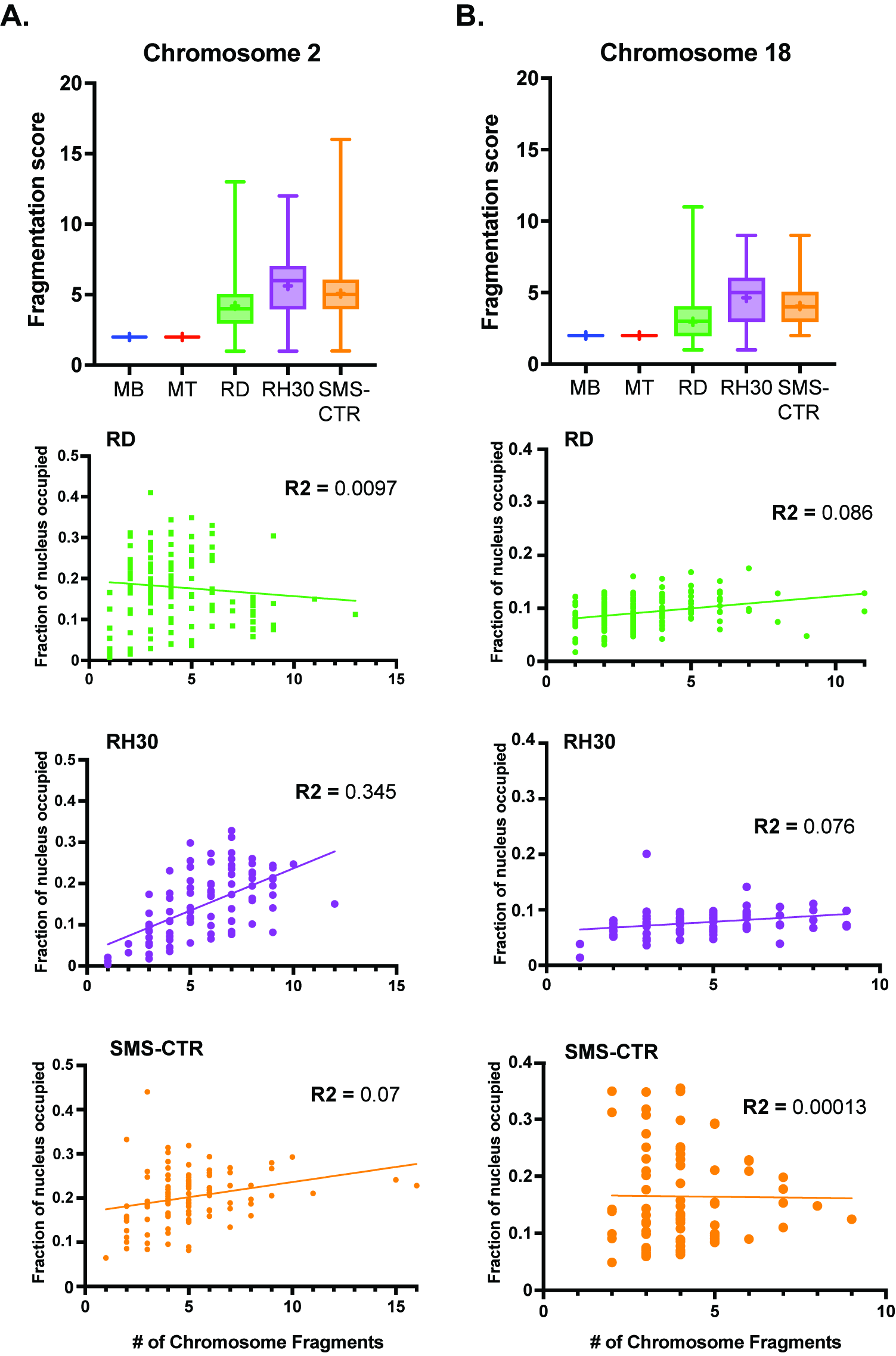

### Supplemental Fig S3

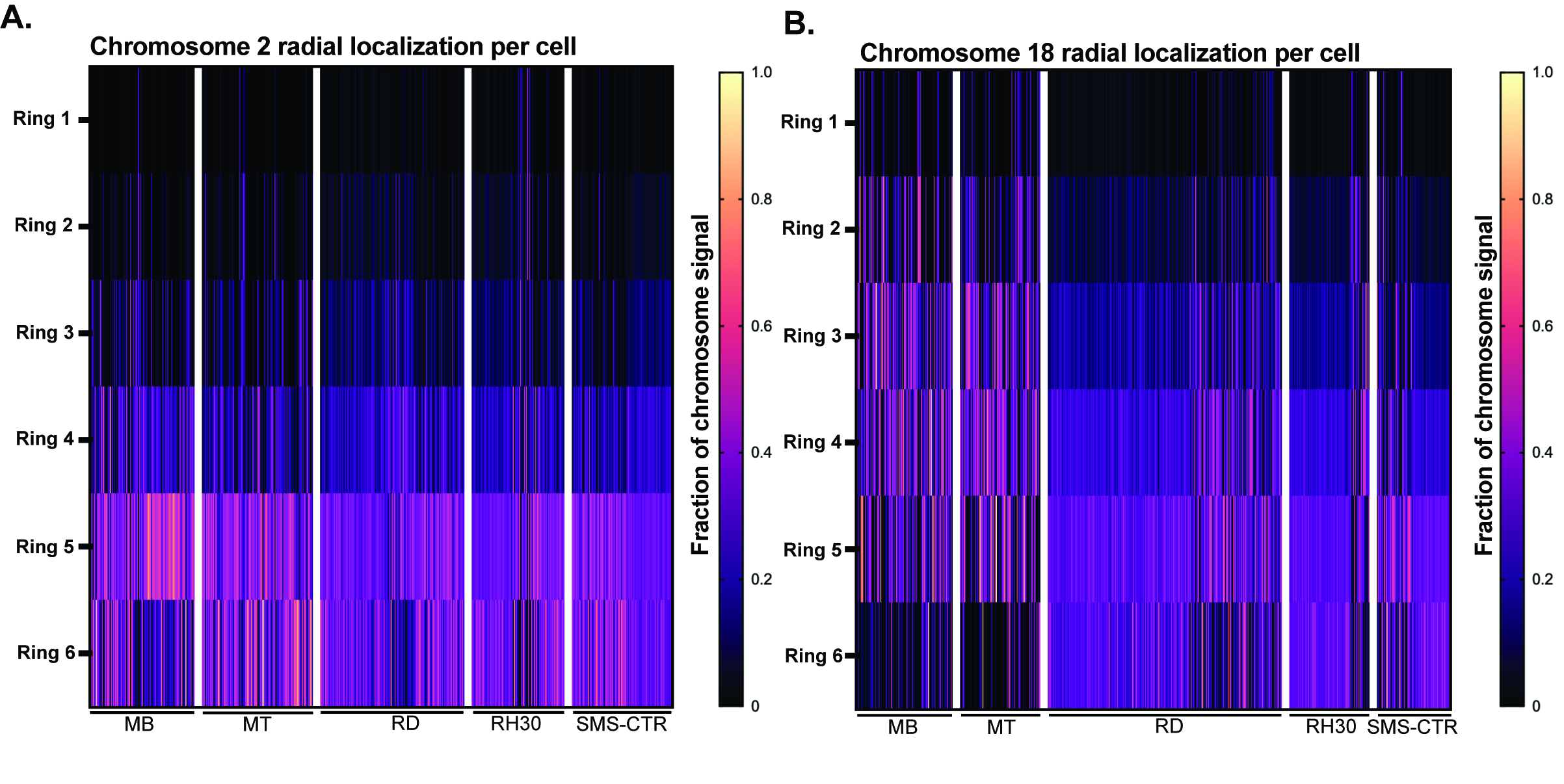

### Supplemental Fig S3

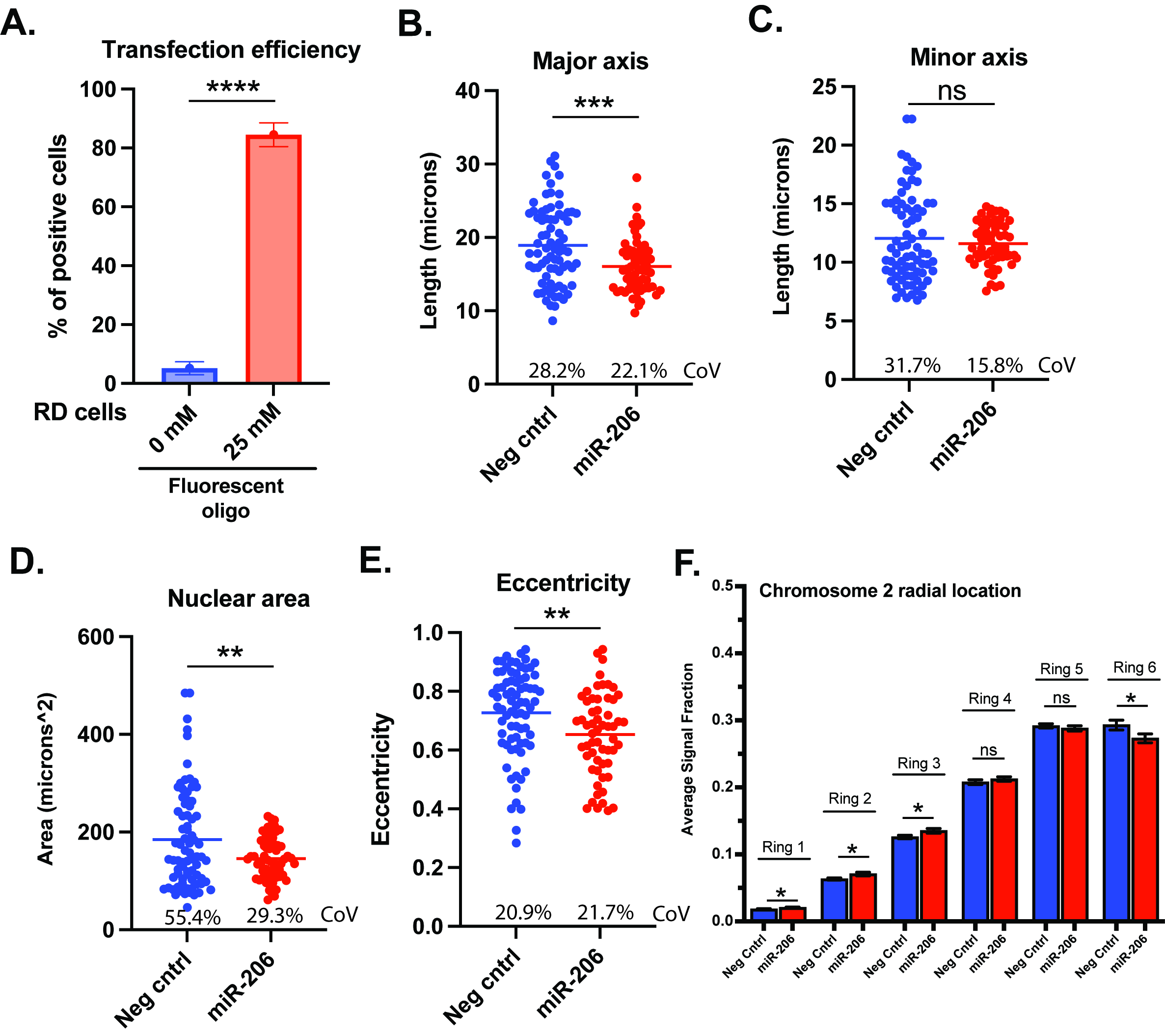
