## Supplemental Table S1 for "Differentiation-dependent chromosomal organization changes in normal myogenic cells are absent in rhabdomyosarcoma cells"

| Cell Type | Coefficient of Variation | | | |
| --- | --- | --- | --- | --- |
|  | **Major axis** | **Minor axis** | **Area** | **Eccentricity** |
| MB | 27.94 | 29.16 | 56.39 | 16.46 |
| MT | 30.22 | 30.30 | 54.38 | 10.08 |
| RD | 32.61 | 39.09 | 66.96 | 18.43 |
| RH30 | 23.97 | 25.49 | 42.91 | 21.03 |
| SMS-CTR | 27.42 | 25.79 | 52.13 | 22.02 |

**Supplemental Table S1**. Coefficients of Variation for Nuclear Characteristic Measurements.
