## Supplemental Table S2 for "Differentiation-dependent chromosomal organization changes in normal myogenic cells are absent in rhabdomyosarcoma cells"

|  |  |  | | Hypothetical | MB | MT | RD | RH30 |
| --- | --- | --- | --- | --- | --- | --- | --- | --- |
|  |  | Major : minor area ratio | | 1:1 | 1.10 : 1 | 1.23 : 1 | 1.17: 1 | 1.17: 1 |
|  |  |  | **Calculations** | | | | | |
| *Analysis Type* | *Chromosomes* | Minor axis calculation | | Minor axis ratio value / (minor axis ratio value + major axis ratio value) | | | | |
|  | *Cells* | Minor axis calculation | | (Minor axis chromosome calculation)^2^ | | | | |
|  |  |  | **Area Adjusted Expected Percentage** | | | | |  |
|  | *Chromosomes* | Major axis | | 50% | 52% | 55% | 54% | 54% |
|  |  | Minor axis | | 50% | 48% | 45% | 46% | 46% |
|  | *Cells* | Major axis | | 75% | 77% | 80% | 79% | 79% |
|  |  | Minor axis | | 25% | 23% | 20% | 21% | 21% |

**Supplemental Table S2.** Expected distribution calculations for chromosome location by cell type.
